# Approximating instantaneous vaccine efficacy from cumulative vaccine efficacy

**DOI:** 10.1101/481804

**Authors:** Kevin van Zandvoort, Andrew Clark, Mark Jit, Stefan Flasche

## Abstract

Few methods exist to estimate vaccine efficacy and its decay following immunisation. Existing methods are largely based on survival analyses such as Poisson or Cox-regression, applied to individual-level data from randomised placebo-controlled trials (RCTs), however, such are often not easily available for analysing pooling evidence across trials. Hence, cumulative vaccine efficacy (VE), the commonly reported endpoint, is implicitly assumed a reasonable proxy for the instantaneous vaccine efficacy (iVE). This assumption is violated if the relative risk (RR) of vaccinated vs unvaccinated is not constant over time, i.e. if vaccine efficacy changes after immunisation. We propose a method to overcome this issue. We use estimates of VE stratified by time since completed immunisation, and estimate time-dependent iVE. We validate the method against simulated data for two forms of vaccine protection: all-or-nothing protection and leaky protection. We illustrate how VE estimates are biased by time-dependent effects in the baseline force of infection and in iVE. Our proposed method improves upon available iVE estimation techniques, particularly if the vaccine induced leaky-like protection and the disease outcome is rare.

## 1 Introduction

Vaccine efficacy is commonly defined as the reduction in the attack rate in vaccinated compared to unvaccinated clinical trial participants, and estimated as 1 – *RR*, where RR is the risk or rate-ratio. If the RR is constant over the period of follow-up for a per protocol analysis (i.e. the time since administration of the final dose in the vaccination schedule), then 1 – *RR* is a reliable indicator of vaccine efficacy, including the instantaneous vaccine efficacy at the end of the follow-up period. However, temporal changes in RR may occur, e.g. due to waning vaccine efficacy after immunisation. Under these circumstances, common estimates of cumulative vaccine efficacy during the time of follow-up (VE) result in biased estimates of instantaneous vaccine efficacy (iVE). Waning of vaccine derived immunity has been reported for most vaccine antigens including mumps[1], pertussis[2], malaria[3], *Streptococcus pneumoniae*[4], and varicella[5].

Several methods have been proposed to estimate iVE. E.g. follow-up time can be stratified in periods, where efficacy is separately estimated within each period, assuming that the time window for each period can be chosen small enough so that iVE remains relatively constant within each[6]. Alternatively, splines can be used to estimate time-dependent hazard rates for survival data[7]. Moreover, Kanaan and Farrington proposed a framework to estimate waning vaccine efficacy, but this requires fitting many parameters and may lead to issues with parameter identifiability[6, 8]. However, many of these methods are based on age- and/or time-dependent parametric survival analysis methods such as Poisson or Cox-regression, which require access to individual-level data. This imposes limitations for pooling vaccine efficacy across studies, as individual-level data is rarely openly shared, in part because of patient identifiability concerns. As a result, trials commonly only report VE alongside average participant follow-up time and imply this to be a good approximation of iVE.

In this manuscript we describe a novel approach for inferring iVE from multiple VE estimates. This is particularly useful in pooled analyses, where 1) pooling multiple survival datasets may be less straightforward, or 2) in meta-regressions, where only the published aggregate estimates are available.

## 2 Estimating vaccine efficacy

### 2.1 Measures of cumulative vaccine efficacy

When calculating VE in clinical trials, the numerator for the risks in intervention and control arms are usually based on cases accrued over the study period, whilst the denominator holds all individuals who were at risk of becoming a case at the start of the study period. This method provides an estimate for the cumulative relative risk, in turn relating to the average vaccine efficacy in the period. However, this is only an accurate estimate of the iVE during the period if such is constant and not e.g. subject to waning.

Similarly, to estimate relative rates, parametric survival analyses such as Poisson or Cox regression often assume that the ratio of hazards in intervention and control arms remain constant with time. Nonparametric methods, such as Kaplan Meier survival estimates, can be used as an alternative[9, 10] to incorporate change in hazard ratios with time as a result of waning of vaccine protection.

While the Kaplan Meier estimates provides an unbiased estimate of VE where the ratio of hazards underlies temporal changes, it provides estimates of VE rather than iVE which are a more intuitive measure of vaccine protection; in particular in the presence of waning vaccine protection VE(t) can be substantially higher than the actual protection at time *t* (iVE(t)).

### 2.2 Approximating instantaneous vaccine efficacy from cumulative vaccine efficacy

Analogue to the Kaplan Meier estimands the probability of not having encountered an infection or disease episode up to time *t* is:

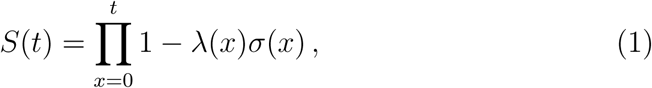

where *λ*(*x*) is the force of infection at time *x* and *σ*(*x*) is the vaccine effect expressed as the instanteneous rate ratio at time x; ie *iV E* = 1 – *σ* and *σ* ≡ 1 in the placebo arm of the study. Hence, cumulative vaccine efficacy can be estimated as via the respective relative risks, *θ*, as:

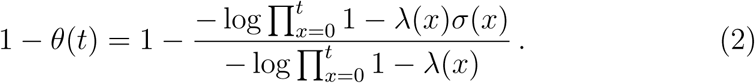

This cannot easily be rewritten to estimate the iVE via *σ*(*x*). However, as the daily force of infection is usually very small (*λ*(*x*) ≪ 1) one can use the Nelson-Aalen estimator and approximate VE(t):

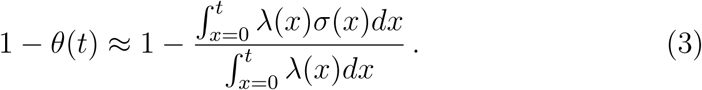

For the special case of proportional hazards that is often assumed in survival analyses, i.e. rate-ratios remain constant over time, equation 3 simplifies to:

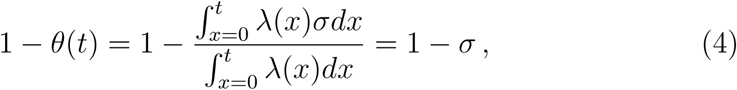

i.e., the cumulative rate-ratio and instantaneous rate-ratio are the same *θ*(*t*) ≡ *σ*. This is irrespective of any change in the baseline-rate *λ*(*x*), assuming that this rate is the same in the vaccinated and unvaccinated groups.

Ideally, infections in the vaccinated and unvaccinated arms are always observed and censored after becoming a case. However, individuals may get asymptomatic infections that are not observed in the study, but boost immunity, especially in the unimmunised individuals. This can alter the baseline rate in the unvaccinated arm of the trial, such that rates in the vaccinated and unvaccinated arm are no longer equivalent. Eventually, this can lead to an overestimation of the relative rate when natural immunity in the unvaccinated arm increases, as has been shown for rotavirus vaccines[11].

Similarly, in the case of waning vaccine-protection, the rate-ratio does not remain constant with time. However, equation 3 can be rewritten as:

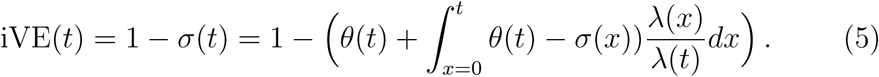

If discretised this provides a recursive equation to calculate iVE(*t*), starting at *σ*(0) = *θ*(0).

## 3 Validation through simulation of an iRCT

### 3.1 Generating simulated data

We simulated data with a deterministic compartmental model (Figure 1). The following differential equations were used to describe the dynamics of infection in an individual randomised controlled trial (iRCT) like setting assuming vaccine protection to act in either an all-or-nothing or leaky fashion. Note that, to simulate an iRCT the force of infection *λ*(*t*) assumes a force of infection that depends on time but not on the number of infected in the trial itself; ie assumes that the force of infection is generated by the surrounding population.

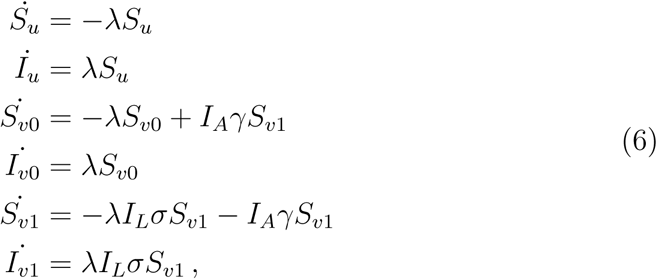

where *S*_*u*_ are the trial participants in the control arm and *S*_*vx*_*, x* ∈ {0, 1} vaccinees who are not immunised or are immunised, respectively (ie for who the vaccine didn’t or did take). For ease of reading, time dependencies are omitted in the equations. It is assumed that all participants were susceptible at time of enrolment. *I*_*u,v*_ then denotes the cumulative number of infections in the control and intervention arms. *I*_*A,L*_ is the indicator function for all or nothing or leaky immunity, respectively:

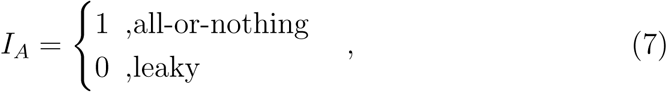

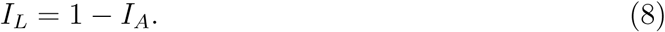

The initial conditions of the model are:

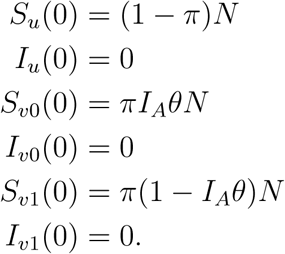

Half (*π* = 0.5) the total number of participants in the trail (*N* = 1000) is randomized to be vaccinated. We further assume that iVE at the start of the trial is 75% (*θ* = 1 – 0.75). We assume waning of vaccine protection to follow a sigmoidal function:

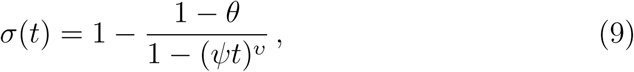

with *ψ* ∈ [1e – 5, 1e – 2] simulating slow or fast waning of vaccine protection, and *v* = 4. In order to assure that iVE is the same in the leaky and all-or-nothing vaccine, we model the rate at which immunized persons lose protection in the all-or-nothing vaccine scenario as:

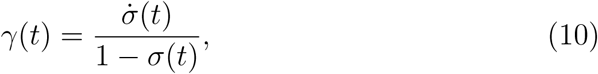

for *σ*(*t*) ∈ [0, 1] (the case *σ*(*t*) = 1, implies *θ* = 1 and hence is not of interest). We further assumed that the force of infection follows a sine-function:

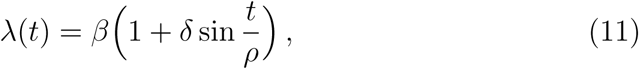

with the baseline force of infection *β* ∈ [2^*-*6^, 2^4^] per 1 000 person days, *δ* ∈ [0.05, 0.4], and *ρ* ∈ [80, 500]. In simulations where the force of infection is assumed non-seasonal, we set *δ* = 0. Table 1 provides an overview of all parameters and their interpretation.

**Figure 1:**
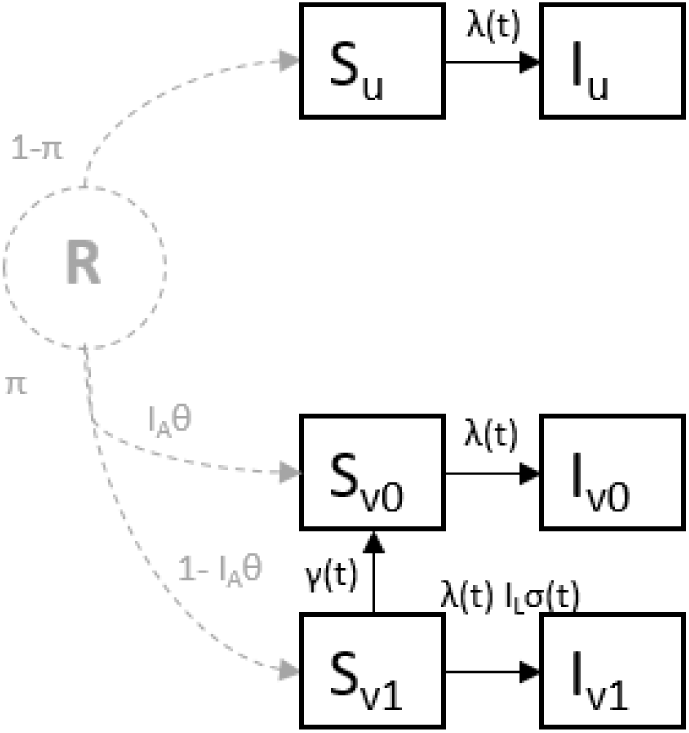
Compartmental model for a hypothetical randomized controlled trial under dierent assumptions of the mode of vaccine action. Compartmental model for the all-or-nothing and leaky mode of vaccine action. Individuals are randomized (R), after which a proportion *π* is vaccinated. All individuals are susceptible at the start of a trial (*S*_*_). Unvaccinated individuals will become infected at rate *λ*(*t*), the force of infection. *I*_*A*_ is 1 for an all-or-nothing vaccine, and 0 otherwise. *I*_*L*_ = 1 *-I*_*A*_. The vaccine will take by a proportion 1 *-I*_*A*_*θ* of the vaccinated individuals (*S*_*v*1_). They will be protected by some degree 1 *-σ*(*t*), where *σ*(*t*) = *I*_*L*_*θ*. In contrast, the vaccine does not take for a proportion *I*_*A*_*θ* (*S*_*v*0_). These individuals become infected at the same rate (*λ*(*t*) as the unvaccinated (*S*_*u*_). In the presence of waning protection, individuals who are fully protected by an all-or-nothing vaccine will lose protection at rate *γ*(*t*). Similarly, the degree of protection that a leaky vaccine offers to vaccinated individuals will wane over time, and *σ*(*t*) will approach 1.

**Table 1:** Parameters in compartmental model, values used in simulations, and their interpretation

| Param | Values | Interpretation |
| --- | --- | --- |
| $N$ | 1000 | Total number of participants in trial. |
| $\pi$ | 0.5 | Proportion of participants that is vaccinated. |
| $I_A$ | $\in \{0, 1\}$ | Variable indicating whether vaccine is all-or-nothing. |
| $I_L$ | $\in \{0, 1\}$ | Variable indicating whether vaccine is leaky. |
| $\lambda(t)$ | $\beta \left( 1 + \delta \sin \frac{t}{\rho} \right)$ | Force-of-infection at time $t$. |
| $\beta$ | $[1\text{e-}3 \cdot 2^{-6}, 1\text{e-}3 \cdot 2^4]$ | Baseline force-of-infection, without seasonal effects. |
| $\delta$ | $\in [0, 0.4]$ | Maximum amplitude of seasonal effect around baseline force-of-infection. |
| $\rho$ | $\in [80, 500]$ | Duration of full seasonal wave is $\rho \cdot 2\pi$ days. |
| $\sigma(t)$ | $1 - \frac{1-\theta}{1-(\psi t)^v}$ | Instantaneous RR at time $t$. iVE can be estimated as $1 - \sigma(t)$. |
| $\gamma(t)$ | $\frac{\dot{\sigma}(t)}{1-\sigma(t)}$ | Rate at which immunized vaccinated individuals lose protection in the all-or-nothing vaccine. Ensures that iVE can be estimated as $1 - \sigma(t)$. |
| $\theta$ | 0.75 | RR after vaccination. VE after vaccination can be estimated as $1 - \theta$. |
| $\psi$ | $\in [1\text{e-}5, 1\text{e-}2]$ | Waning rate. The half-life of protective antibodies is $1/\psi$ days (days at which $iVE = 1 - \frac{2-\theta}{2}$). |
| $v$ | 4 | Controls the shape of the waning curve. $\sigma(t)$ will have a sigmoidal shape when $v > 1$. When $v \rightarrow \infty$, $\sigma(t)$ becomes a stepfunction. |

### 3.2 Interpreting simulated iVE

We ran the models for 2000 days. Models were set up in such a way that the assumed iVE in each simulation can be calculated as 1 *-σ*(*t*) for both all-or-nothing and leaky vaccines. Do note that iVE should be derived using a measure of the instantaneous risk ratio for all-or-nothing vaccines, whilst it should be derived using a measure of the instantaneous rate ratio for leaky vaccines. An alternative, more intuitive way to calculate instantaneous rate ratios is:

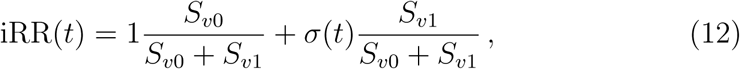

which is a weighted average of the rate ratio in the vaccinated individuals who are still susceptible. It reduces to *iRR*(*t*) = *S*_*v*0_*/*(*S*_*v*0_ + *S*_*v*1_) for the all-or-nothing vaccine and to *iRR*(*t*) = *σ*(*t*) for the leaky vaccine. Similarly, we can calculate instantaneous risk-ratios as:

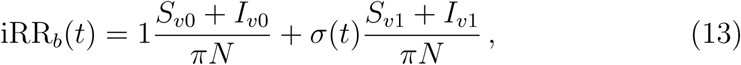

which is a weighted average of the risk ratio in all vaccinated individuals, not censoring those who were infected. This reduces to 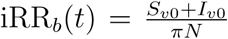 (the proportion vaccinated that lost immunity) in the all-or-nothing vaccine, and to iRR_*b*_(*t*) = *σ*(*t*) in the leaky vaccine. Note that iRR(*t*) = iRR_*b*_(*t*) for the leaky vaccine.

Whilst these instantaneous measures will not be observed in iRCTs, cumulative case-counts are. We used the cumulative case-counts of our simulated trials to estimate VE twice, once using risk ratios, using the conventional formula (risk in vaccinated divided by risk in unvaccinated), and once using a ratio of cumulative Kaplan-Meier hazard-estimates. We then applied our method, given in equation 5, to convert the ratios of cumulative Kaplan-Meier hazard-estimates to instantaneous rate ratios. This was done twice, once controlling for the simulated seasonal effects (the 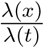 component in equation 5), and once assuming that there were no seasonal effects, as this will often be a necessary constraint in practical applications.

Ultimately, we compared these estimated measures of iVE to the assumed iVE (1 – *σ*(*t*)) in each simulation, to validate how our method performed under different conditions. Incididence rates were varied across simulations. The preventable outcome of interest is common in simulations where the incidence rate is high, whereas it is rare when the incidence rate is low.

## 4. Analysis of simulated data

### 4.1 Assumed instantaneous measures

Figure 2 shows a subset of our results, where the first two columns show the assumed iVE calculated as 1 – *iRR*_*b*_ and 1 – *iRR*. These measures of iVE are usually not directly observed in a trial. Coincidentally, as shown before, 1 – *iRR*_*b*_ in the first column equals to 1 – *σ*(*t*) for both all-or-nothing and leaky vaccines.

**Figure 2:**
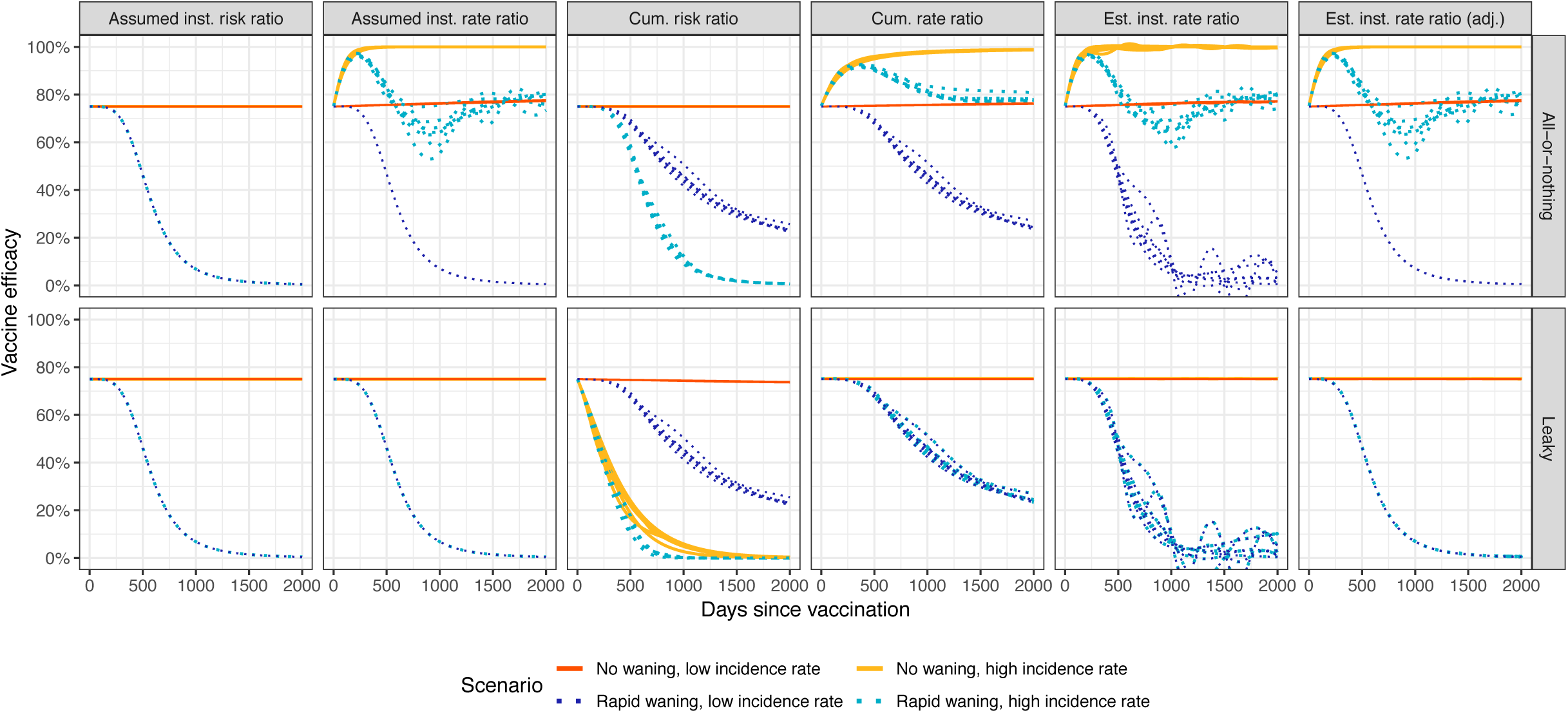
Subset of results for simulations with and without waning vaccine efficacy. The top row shows results for the all-or-nothing vaccine, whilst the bottom row shows results for the leaky vaccine. Each column represents vaccine-efficacy as derived using a different method. From left to right, this is 1) the assumed iVE calculated as 1 – *iRR*_*b*_ (which is the correct iVE for the all-or-nothing vaccine); 2) the assumed iVE calculated as 1 – *iRR* (which is the correct iVE for the leaky vaccine); 3) the VE derived from risk ratios; 4) the VE derived from cumulative hazard ratios; 5) the estimated iVE, assuming no seasonal effects; and 6) the estimated iVE, adjusting for known seasonal effects. Solid lines represent scenarios where vaccine efficacy does not wane. Their scenarios with a low incidence rate are shown in orange, and scenarios with a high incidence rate are shown in yellow. Dotted lines represent scenarios where vaccine efficacy does wane over time. Their scenarios with a low incidence rate are shown in dark-blue, whereas scenarios with a high incidence rate are shown in light-blue. The different lines within each scenario represent different seasonal effects.

In simulations where iVE does not wane, shown by the red and yellow lines, iVE remains at 75% for both modes of vaccine action. However, when using the instantaneous rate ratio to estimate iVE for an all-or-nothing vaccine, it will appear to wax over time. This is explained by equation 12, which shows that this measure of iVE is a weighted average of the iVE in those vaccinated individuals who are still susceptible. Assuming that time approaches ∞ and the force-of-infection remains the same, ultimately all vaccinated but unprotected individuals become infected. The weighted average of iVE is then dominated by those who are immunized and fully protected, and hence, *iV E* → 100%. Moreover, the pool of vaccinated but unprotected susceptibles depletes faster when the incidence rate is high (illustrated by the yellow line). Therefore, this measure of iVE approaches 100% much faster in simulations with high incidence rates. In all simulations shown in figure 2, *β* = 1e−3·2^−4^ and *β* = 1e−3·2^3.5^ are used in simulations with low and high incidence rates, respectively.

In simulations where iVE does wane (shown by the dark- and light-blue lines), iVE starts at 75% and diminishes towards 0% as immunity is lost. In all with-waning simulations shown in figure 2, *ψ* = 1e – 2.75, so half of iVE will have waned after about 560 days (*iV E* = 37.5%).

Again the iVEs as estimated by 1 – *iRR*_*b*_ and 1 – *iRR* give the correct measures of 1 – *σ*(*t*) for the all-or-nothing vaccine and the leaky vaccine, respectively. Note that, for the all-or-nothing vaccine, iVE derived from the relative risk is similar to that derived from the relative rate when the incidence rate is low, especially when iVE wanes rapidly. This is due to 1) the well-known fact that risks and rates are numerically similar when an outcome is rare; and 2) the fact that, in this scenario, the rate at which those vaccinated and fully protected lose immunity is stronger than the rate at which those vaccinated but unprotected become infected.

The opposite happens when the incidence rate is high, here shown by the light-blue lines. Although this measure of iVE provides an estimate of the level of protection in those vaccinated and (still) susceptible, comparison to the first column shows that this is not a good proxy for the iVE for all vaccinated individuals when the incidence rate is high. Note that the light-blue lines of the different simulations do not overlap, indicating that this estimate of iVE is affected by seasonal effects.

### 4.2 Observable measures of VE

Whereas the iVE measures discussed in the previous section are not observed in a trial, cumulative case-counts are. It are estimates of VE using these cumulative counts that are usually reported in RCTs. In figure 2, VEs derived from the observed risk ratio are shown in the third column, whereas VEs derived from the observed cumulative hazard ratios are shown in the fourth column.

First, for the all-or-nothing vaccine, in simulations without waning, the measure of VE derived from the risk ratio is an appropriate measure of iVE. However, measures of VE derived from cumulative hazard ratios will approach 100% for the same reason as happens for those of iVE derived from 1 *-iRR*_*b*_.

Second, for the leaky vaccine, in the absence of waning, the measure of VE derived from cumulative hazard ratios is an appropriate measure of iVE. However, measures of VE derived from risk ratios will approach 0%, which is especially obvious when the incidence rate is high. This occurs because infected individuals are not censored when estimating a risk ratio. As leaky vaccines do not provide 100% protection, assuming that the force of infection remains constant and time approaches ∞, all vaccinated individuals will eventually become infected. Thereby, the risks in the vaccinated and unvaccinated strata converge, and *V E* → 0%. A more detailed overview of these effects is given by Smith et al[12].

When iVE does wane, both VE derived from a risk ratio for an all-or-nothing vaccine and VE derived from a cumulative hazard ratio for a leaky vaccine will overestimate the actual iVE. These measures are both influenced by historic values of VE. Therefore, VE at time t provides an estimate of the average iVE up until time t, rather than the iVE at time t. As a result of this averaging behaviour, seasonal effects barely influence these estimates. Again, for both all-or-nothing and leaky vaccines, VEs derived from risk and rate ratios are similar when the incidence rate is low.

### 4.3 Estimated iVE

We apply the method in equation 5 to estimate iVE by converting the observed measures of the cumulative hazard ratios, shown in the fourth column of figure 2. We do this once by assuming no seasonal effects, shown in the fifth column, and once by adjusting for seasonal effects, shown in the sixth column. Note that adjusting for seasonal effects requires knowledge about the force of infection over the entire period. This will not be available in most practical applications, but can be done here as data is simulated.

When adjusting for seasonal effects, the estimated iVEs retrieve the assumed iVEs derived from 1 – *iRR* perfectly, for both all-or-nothing and leaky vaccines. As this is not the iVE one is usually interested in for all-or-nothing vaccines, the appropriateness of this method for all-or-nothing vaccines will depend on the similarity between the risk and rate ratio, which in turn is influenced by the incidence rate of the outcome of interest.

When not adjusting for seasonal effects, as in the fifth column, the deviation between the estimated and assumed iVE will depend on the extent at which seasonality affects the incidence rate. The unadjusted iVE estimates clearly show the sine-functions used in simulating seasonal effects, whereas these effects were barely noticeable in measures of VE. Do note however, that even these unadjusted measures provide better estimates of iVE when compared to VE based on cumulative counts.

### 4.4 Comparing estimated iVE to assumed iVE

To more formally identify instances where this method fails or works, we plotted the maximum percentage point (pp) deviation at any day of the trial between the estimated iVE and the actual assumed iVE in figure 3. Simulations use multiple assumptions about the incidence rate (y-axes), and the rate at which immunity wanes (x-axes).

**Figure 3:**
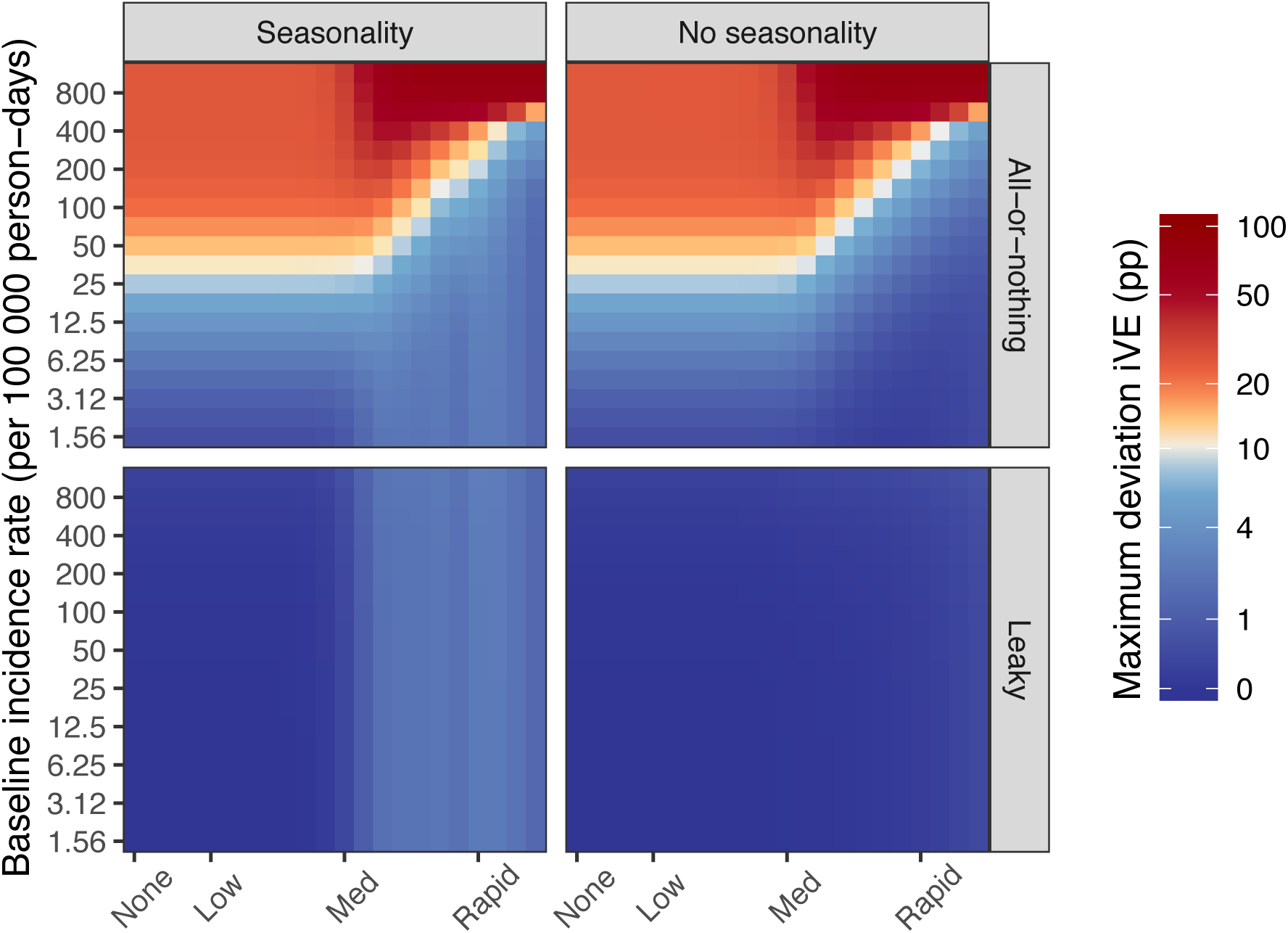
Maximum deviation in percentage point (pp) between the assumed and estimated unadjusted iVE. Maximum deviation in percentage point (pp) at any time between the assumed and estimated unadjusted iVE. The top row shows results for the all-or-nothing vaccine, whilst the bottom row shows results for the leaky vaccine. The first column shows results for simulations with seasonal effects, whilst seasonal effects were not used in the right column. The y-axes represent scenarios with different incidence rates, whilst the x-axes represent scenarios with different degrees at which iVE wanes.

All estimated iVE values were generated assuming no seasonal effects. However, results in the right column of figure 3 would look exactly the same to those in left column if we would have adjusted for those effects. In simulations with seasonality, seasonal effects are modelled using *δ* = 0.05 and *ρ* = 80.

Although deviations are large when applied to an all-or-nothing vaccine and when the incidence rate is high, this deviation becomes much smaller when the outcome becomes less common. Even with moderate values of the incidence rate, the deviation is relatively small when immunity wanes rapidly.

For leaky vaccines, iVE is barely effected by the incidence rate, which is expected. When no seasonal effects are simulated, this method is always able to retrieve the actual assumed iVE for leaky vaccines, as the deviation is 0 pp. However, when we cannot adjust for seasonality, the estimated iVE will somewhat deviate from the actual iVE. Although this deviation is not related to the incidence rate, it becomes larger when immunity wanes more rapidly.

Sensitivity analyses with different values of *δ* and *ρ* are provided in Appendix A. When seasonal effects are stronger, the deviation between the estimated and assumed iVE may be significant when these effects are not accounted for. Due to the interplay between waning and seasonal effects, this deviation will be higher when immunity wanes more rapidly.

## 5 Limitations

There are several limitations to the method presented here. First, although it can effectively retrieve estimates of iVE based on instantaneous rate ratios, this is not the measure of iVE usually of interest for all-or-nothing vaccines. Therefore, the appropriateness of this method for all-or-nothing vaccines will depend on how common the preventable outcome of interest is.

Second, when strong seasonal effects occur, which is common for some infectious agents, these changes in the force of infection will interact with any waning of immunity. Ideally, these seasonal effects should be taken into account when converting VE. Although one may fit a natural history model to estimate such effects, they will often be unknown. Then, the estimated iVE will only approximate the true iVE, whereby the acceptable magnitude of the error is likely disease and outcome specific.

Third, this method requires an iterative approach to convert values of VE to iVE. Therefore, detailed time-specific knowledge of VE is required. Successive steps used in this iterative approach should be small enough in order to accurately capture the extent of waning.

Fourth, this method assumes that the true VE is known, whereas there will always be some level of uncertainty around estimates of VE. One may use bootstrap sampling to reflect this uncertainty in converting VE to iVE. The mean and/or median of the iVEs in all bootstrap samples may then be used as the central estimate of iVE, whilst their quantiles can be used to report the level of confidence required. We did not investigate the effects of underreporting of the outcome variable on estimates of iVE, as this would inheritely bias estimates of VE as well.

Lastly, we applied this method to simulations for all-or-nothing and leaky vaccines. We acknowledge that methods of vaccine action may take different forms, such as combinations of these two extremes, or that individual immunity may be much more heterogeneous in practice. However, many other existing methods assume either all-or-nothing or leaky effects, and few investigate heterogeneous vaccine responses.

## 6 Concluding remarks

Despite its limitations, we have shown that this method is effective in converting cumulative VEs to iVEs for leaky vaccines, and may also be useful for all-or-nothing vaccines, in instances where the outcome is rare.

There can be many challenges in studying VE, including the effect of asymptomatic infections boosting immunity, heterogeneities in vaccine responses, and imperfect biological markers. This method may assist researchers with a new way to investigate waning of iVE.

Ideally, approximated iVEs derived using this method should be compared to true iVEs: immunological markers which provide a good correlate or surrogate of protection. Furthermore, the effect of the relationship between seasonality and waning iVE deserves more attention in future studies.

## Supporting information

## Acknowledgements

This work was supported by the Bill & Melinda Gates Foundation (BMGF). Kevin van Zandvoort, Andrew Clark, and Mark Jit are supported by the Vaccine Impact Modelling Consortium, jointly funded by Gavi, the Vaccine Alliance and the Bill & Melinda Gates Foundation. Stefan Flasche is supported by a Sir Henry Dale Fellowship jointly funded by the Wellcome Trust and the Royal Society (Grant number 208812/Z/17/Z).

