## Supplementary material for "Approximating instantaneous vaccine efficacy from cumulative vaccine efficacy"

### Appendix A. Exploring the effect of different assumptions about seasonality on our results

We performed a sensitivity analysis to look at different assumptions about seasonality on our results. In figure 2 in our main paper, we showed the results where the amplitude of the seasonal patterns was $\delta = 0.05$ (i.e. a maximum 5% in- or decrease in the baseline rate) and $\rho = 80$ (i.e. very short seasonal waves). The plots in figure 2 are therefore the same as those shown in the top-left corner of figures A.1 and A.3 for the all-or-nothing vaccine, and figures A.2 and A.4 for the leaky vaccine. Whilst figure 2 showed the results for a fixed incidence rate, these are the same as the results of simulations with a change in the incidence rate, where these changes are adjusted for (as in figures A.3 and A.4). For the same reason, values in the plots in figures A.3 and A.4 are exactly the same across the grid, regardless of the extent of seasonality in the force-of-infection.

In all plots in this appendix, different rows represent different durations of seasonal patterns, where lower numbers lead to short wavelengths and higher numbers lead to long wavelengths. Moreover, different columns represent the different amplitudes that the force-of-infection can be amplified with (i.e. 5, 10, 20, or 40% higher or lower). Therefore, the extent of seasonality becomes higher when moving from left to right in the grid, and when moving from bottom to top in the grid.

When seasonal patterns in the baseline rate become stronger, the deviation between the approximated and real iVE will significantly increase if not accounted for. Due to the interplay between waning and potential changes in this rate, this deviation will be higher as iVE wanes more rapidly for leaky vaccines, whilst it will be lower when iVE wanes more rapidly for all-or-nothing vaccines.

#### Maximum deviation in percentage point (pp) between the real and approximated unadjusted iVE for all-or-nothing vaccines

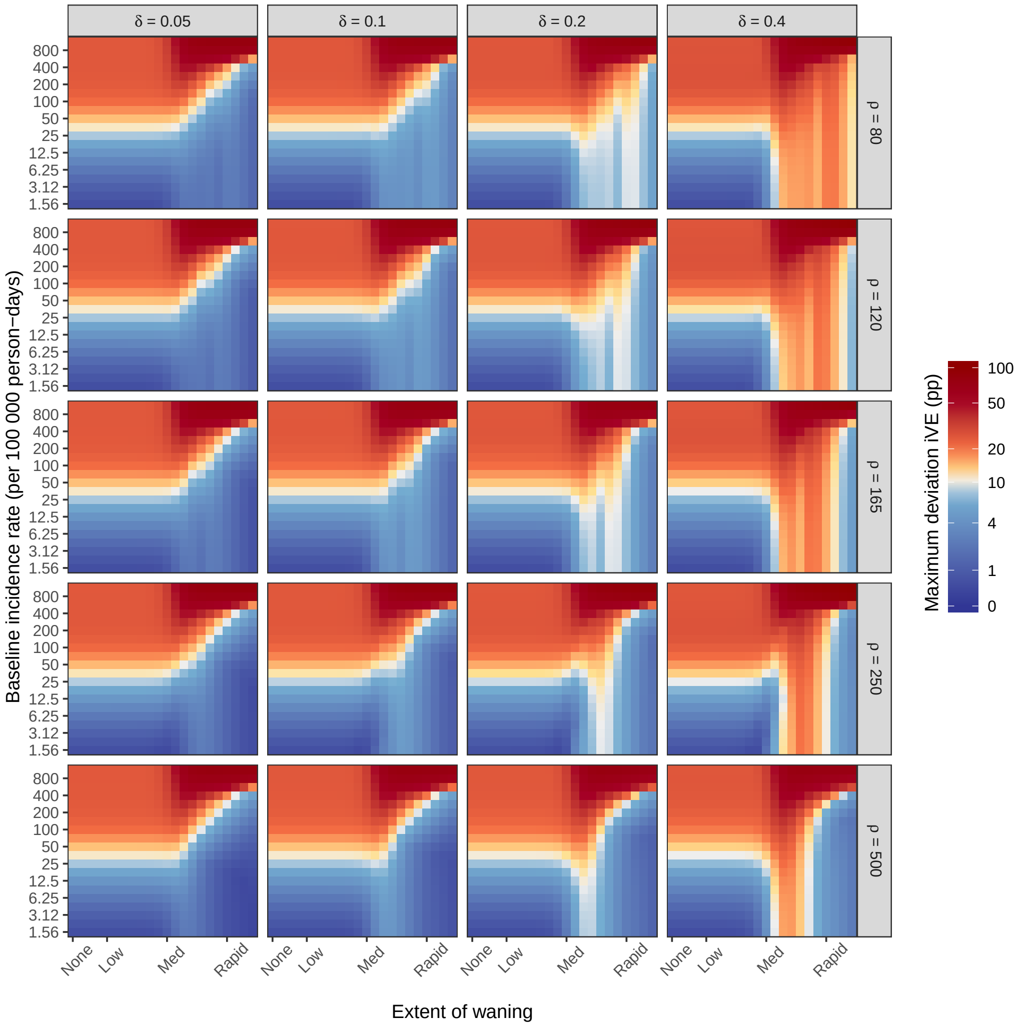


Figure A.1: The maximum percentage point (pp) deviation between the true instantaneous vaccine efficacy (iVE) and the approximated iVE (unadjusted for seasonal patterns in the force-of-infection) for an all-or-nothing vaccine. Approximated iVEs are compared against the true iVE derived from instantaneous risk ratios. Different rows represent different durations of seasonal patterns, where lower numbers lead to short wavelengths and higher numbers lead to long wavelengths. Different columns represent the different amplitudes that the force-of-infection can be amplified with (i.e. 5, 10, 20, or 40% higher or lower). The extent of seasonality becomes higher when moving from left to right in the grid, and when moving from bottom to top in the grid. The plot in the top-left corner ($\delta = 0.05$, $\rho = 80$) is presented in figure 2 in the main body of this paper.

**Maximum deviation in percentage point (pp) between the real and approximated unadjusted iVE for leaky vaccines**

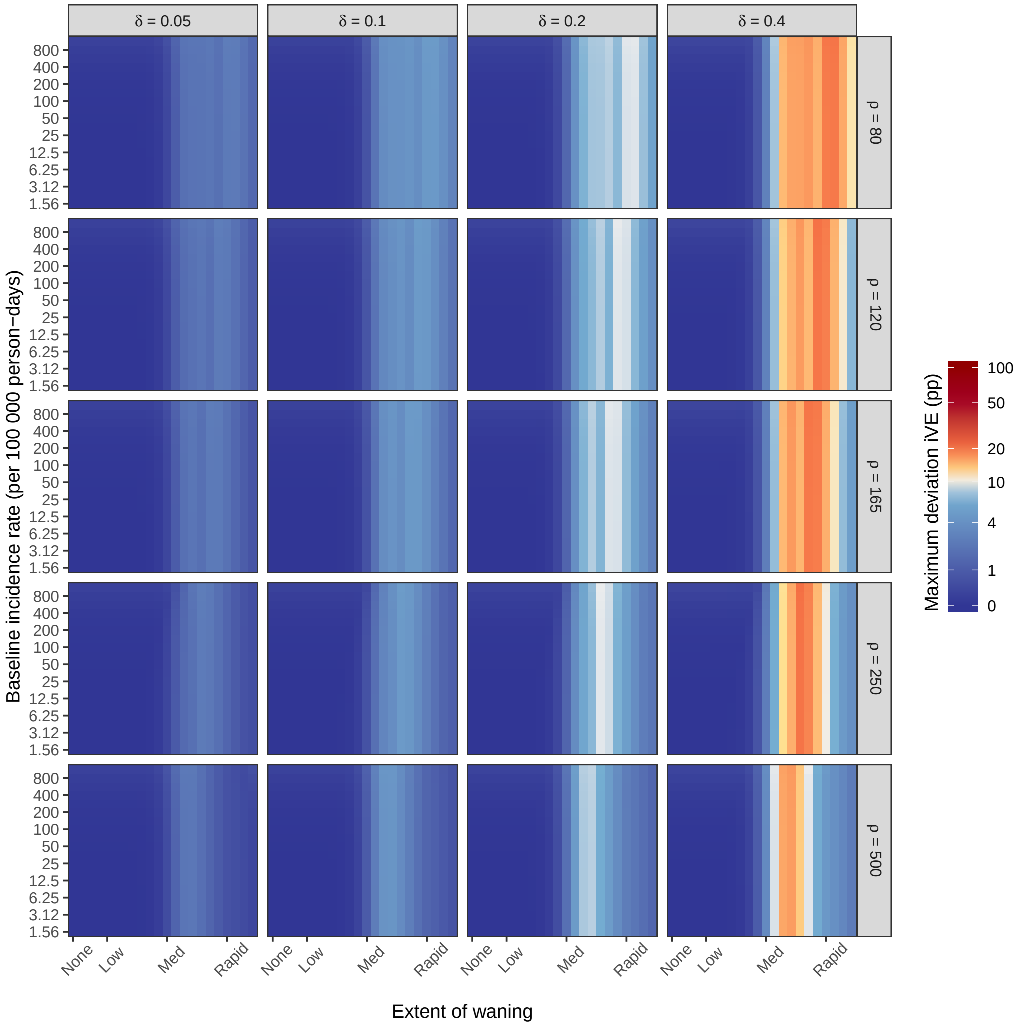


Figure A.2: The maximum percentage point (pp) deviation between the true instantaneous vaccine efficacy (iVE) and the approximated iVE (unadjusted for seasonal patterns in the force-of-infection) for a leaky vaccine. Approximated iVEs are compared against the true iVE derived from instantaneous rate ratios. Different rows represent different durations of seasonal patterns, where lower numbers lead to short wavelengths and higher numbers lead to long wavelengths. Different columns represent the different amplitudes that the force-of-infection can be amplified with (i.e. 5, 10, 20, or 40% higher or lower). The extent of seasonality becomes higher when moving from left to right in the grid, and when moving from bottom to top in the grid. The plot in the top-left corner ($\delta = 0.05$, $\rho = 80$) is presented in figure 2 in the main body of this paper.

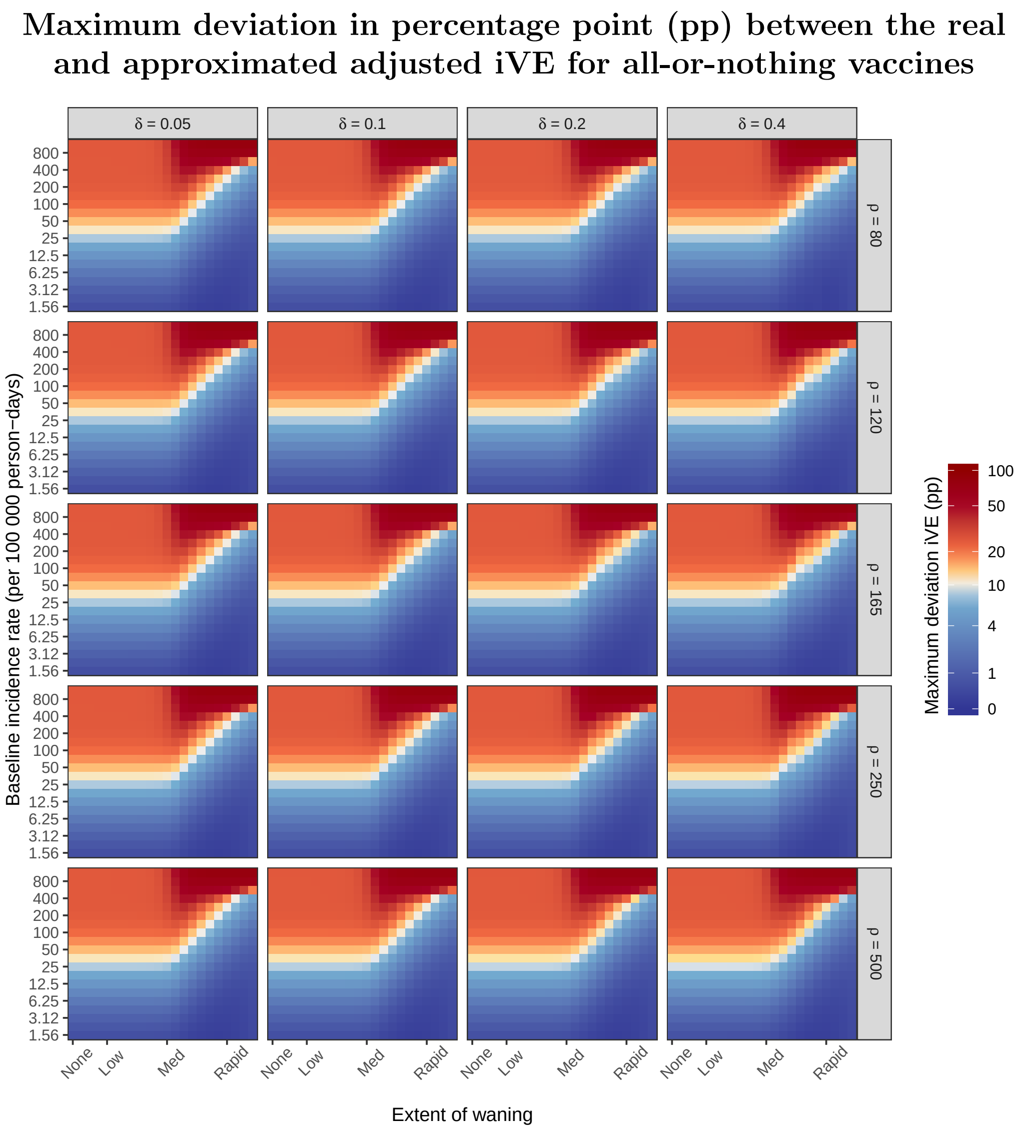


Figure A.3: The maximum percentage point (pp) deviation between the true instantaneous vaccine efficacy (iVE) and the approximated iVE (unadjusted for seasonal patterns in the force-of-infection) for an all-or-nothing vaccine, when adjusting for changes in the force-of-infection. Approximated iVEs are compared against the true iVE derived from instantaneous risk ratios. Different rows represent different durations of seasonal patterns, where lower numbers lead to short wavelengths and higher numbers lead to long wavelengths. Different columns represent the different amplitudes that the force-of-infection can be amplified with (i.e. 5, 10, 20, or 40% higher or lower). The extent of seasonality becomes higher when moving from left to right in the grid, and when moving from bottom to top in the grid. The influence of seasonality disappears when adjusted for, and therefore all plots in this grid are the same. They are the same as the plot presented in the "Fixed incidence rate" column for the all-or-nothing vaccine in figure 2 of our main paper.

#### Maximum deviation in percentage point (pp) between the real and approximated adjusted iVE for leaky vaccines

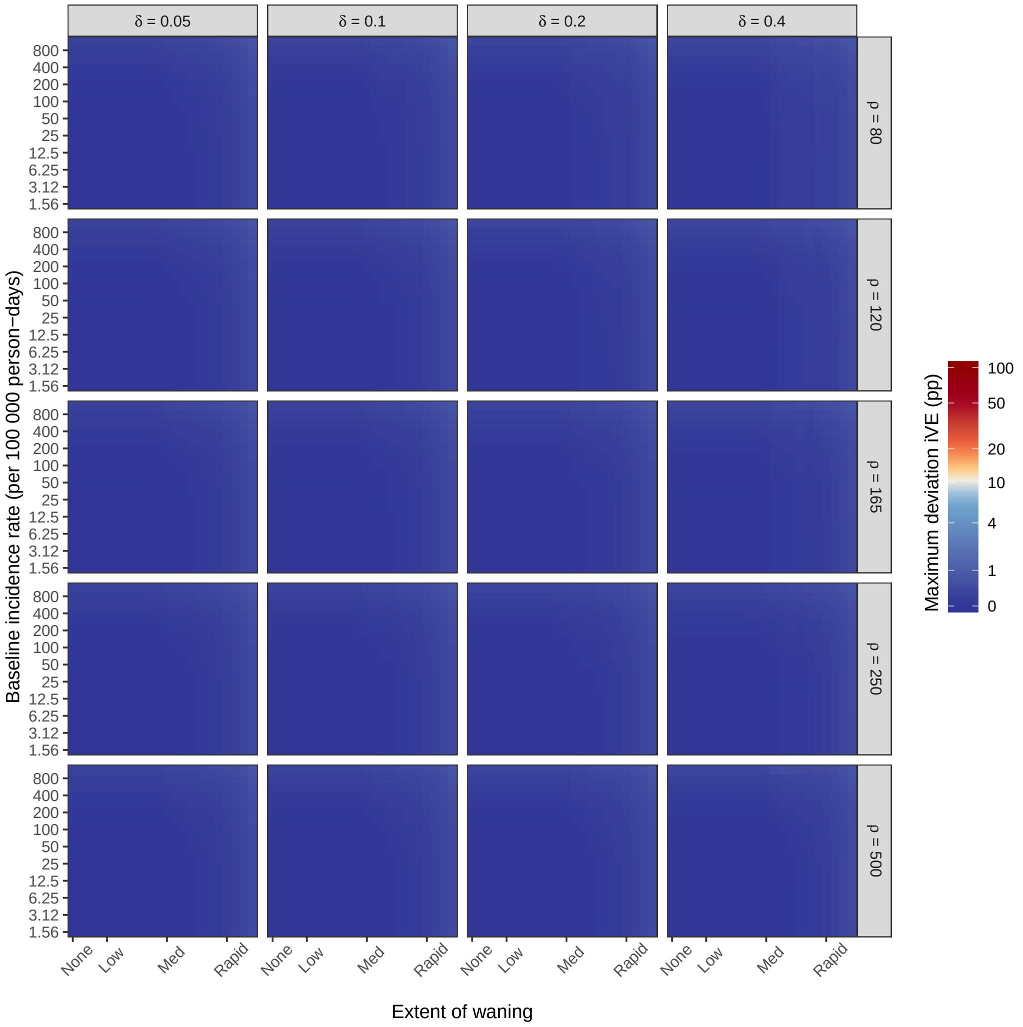


Figure A.4: The maximum percentage point (pp) deviation between the true instantaneous vaccine efficacy (iVE) and the approximated iVE (unadjusted for seasonal patterns in the force-of-infection) for a leaky vaccine, when adjusting for changes in the force-of-infection. Approximated iVEs are compared against the true iVE derived from instantaneous rate ratios. Different rows represent different durations of seasonal patterns, where lower numbers lead to short wavelengths and higher numbers lead to long wavelengths. Different columns represent the different amplitudes that the force-of-infection can be amplified with (i.e. 5, 10, 20, or 40% higher or lower). The influence of seasonality disappears when adjusted for, and therefore all plots in this grid are the same. They are the same as the plot presented in the "Fixed incidence rate" column for the leaky vaccine in figure 2 of our main paper.
